# Antero-posterior versus lateral vestibular input processing in human visual cortex

**DOI:** 10.1101/530808

**Authors:** Felipe Aedo-Jury, Benoit R. Cottereau, Simona Celebrini, Alexandra Séverac Cauquil

## Abstract

Visuo-vestibular integration is crucial for locomotion, yet cortical mechanisms involved remain poorly understood. We combined binaural monopolar galvanic vestibular stimulation (GVS) and functional magnetic resonance imaging (fMRI) to characterize the cortical networks activated during antero-posterior and lateral stimulations in humans. We focused on functional areas that selectively respond to egomotion-consistent optic flow patterns: the human middle temporal complex (hMT+), V6, the ventral intraparietal (VIP) area, the cingulate sulcus visual (CSv) area and the posterior insular cortex (PIC). Areas hMT+, CSv, and PIC were equivalently responsive during lateral and antero-posterior GVS while areas VIP and V6 were highly activated during antero-posterior GVS but remained silent during lateral GVS. Using psychophysiological interaction (PPI) analyses, we confirmed that a cortical network including areas V6 and VIP is engaged during antero-posterior GVS. Our results suggest that V6 and VIP play a specific role in processing multisensory signals specific to locomotion during navigation.

## Introduction

Self-motion (egomotion) perception permits us to estimate our on-going change of position within the surrounding space in order to properly interact with our environment. In the brain, egomotion is processed from multisensory inputs, particularly vestibular and visual ones whose integration remains poorly understood.

In macaques, several groups have shown vestibular projections in the medial superior temporal area (MST), a visual area involved in motion and self-motion perception based on optic flow (Duffy, 1998; Bremmer et al., 1999; Gu et al., 2006). MST projects towards the ventral intraparietal area (VIP) that is sensitive to visual heading and receives vestibular inputs (Klam and Graf, 2003a, b). A recent study demonstrated that the visual posterior area (VPS), an area located at the posterior end of the sylvian fissure, also contains multi-sensory neurons that process both optic flow and vestibular signals (Chen A et al., 2011).

In humans, neuroimaging studies revealed several brain regions involved in visual egomotion processing. For example, Wall and Smith (2008) found that the ventral intraparietal (VIP) and the cingulate sulcus visual (CSv) areas had selective responses to optic flow patterns that are compatible with those that receive our retina during locomotion (i.e. a selectivity to egomotion-consistent optic flows). A preference for egomotion-consistent visual pattern, although weaker, was also reported in human MST, (Morrone et al., 2000). Human MST might therefore constitute an intermediate stage of egomotion processing which is further developed in areas VIP and CSv. A follow-up study (Cardin and Smith, 2010) used wide-field visual stimuli to demonstrate that putative area V6 and two vestibular areas were also included in a network processing egomotion. Using galvanic vestibular stimulation (GVS), Smith et al., (2012, see also Billington Smith, 2015) showed that MST and CSv were also vestibularly-driven, which strengthen their role in egomotion processing. In the same study, responses to GVS in V6 and VIP were very weak if not absent. However, these authors used the classical binaural bipolar configuration where the anode is placed on one mastoid and the cathode on the other. In this case, GVS is known to elicit a lateral postural tilt towards the anode when the body is free to move, (e.g., Njiokiktjien and Folkerts, 1971, Nashner and Wolfson, 1974, Lund and Broberg, 1983), but also a feeling of motion in the opposite, yet lateral, direction (Fitzpatrick et al., 2002; and Day, 2004). These responses are compatible with an activation of the parts of the vestibular apparatus sensitive to roll tilt, in the frontal plane (Day BL, et al., 1997; Séverac Cauquil et al., 1993). Therefore, such a GVS design prohibits the investigation of the contribution of antero-posterior motion signal. Yet, human motion, in particular locomotion, mostly refers to forward displacements: it principally includes translational egomotion in the postero-anterior (i.e., forward) direction. If walking and running involve a complex pattern of acceleration and deceleration that also comprises vertical translation and sagittal rotation, these components are nevertheless minimized in order to stabilize the head (Pozzo et al., 1990). The cortical networks engaged in visuo-vestibular integration during antero-posterior egomotion might therefore be different from those involved during lateral egomotion. The different pathways followed for motion in depth processing compared to lateral motion processing supports this hypothesis (Cottereau et al., 2014). So does the finding that different areas such as MST and V6 would encompass dissociated components of self-motion from optic flows, i.e., heading for the former and obstacle avoiding for the latter (Cardin et al., 2012a).

In the present study, we (1) reproduced Smith’s et al paradigm using different stimulation parameters so as to determine whether the set of visual areas that they found can be reliably activated by a different type of lateral GVS. We also (2) determined whether antero-posterior vestibular inputs activated a different cortical network. In that aim, we used binaural monopolar GVS, since this design, although much less usual, has been shown to induce a body response and self-motion illusion in the antero-posterior plane (Magnusson et al., 1990; Séverac Cauquil A et al., 1998; 2000; Aoyama et al., 2015): forward when anodes are on the forehead and backward with anodes over the mastoid processes. Such postural tilts in the antero-posterior direction fit with Day et al’s model (2011). They postulate that by polarizing equally both vestibular apparatus, binaural monopolar GVS provides a fake backward or forward self-motion input. Among several studies indicating that GVS induces a postural tilt towards the anodes, counteracting the vection direction (away from the anodes), the most recent demonstrated the perfect adequacy between subjective perceptual responses and objective quantified head movements, for both lateral and antero-posterior GVS stimulations (Aoyama et al., 2015). Here, we combine this tool with fMRI to differentiate the visual cortical networks activated during antero-posterior (AP) and lateral, (L) GVS.

## Materials & Methods

### Participants

Thirteen healthy human subjects (mean age 28.4, range 19-45, 7 females) were included in this study. Eleven were right handed, as assessed with the Edinburgh Inventory (Oldfield, 1971). They all participated in the galvanic stimulation experiment. 11 of them also performed an additional experiment that included functional localizers. All subjects had normal or corrected-to-normal vision, reported no history of neurological or psychiatric disease, and gave written informed consent before participation, in accordance with the Declaration of Helsinki. This study was approved by the local ethic committee (ID RCB: 2012-A01052-41). Subjects received 80 euros of monetary compensation for their participation.

### Stimuli and design

#### Galvanic stimulation

Vestibular stimuli consisting of 2s of 1mA square-pulses were delivered by two identical dedicated current-limited stimulators (DS5, Digitimer, UK, CE certified for biomedical research N(IEC) 60601) through 4 disposable carbon electrodes (Skintact, FSWB00) placed on the forehead and over the mastoid processes (see Figure 1).

**Figure 1:**
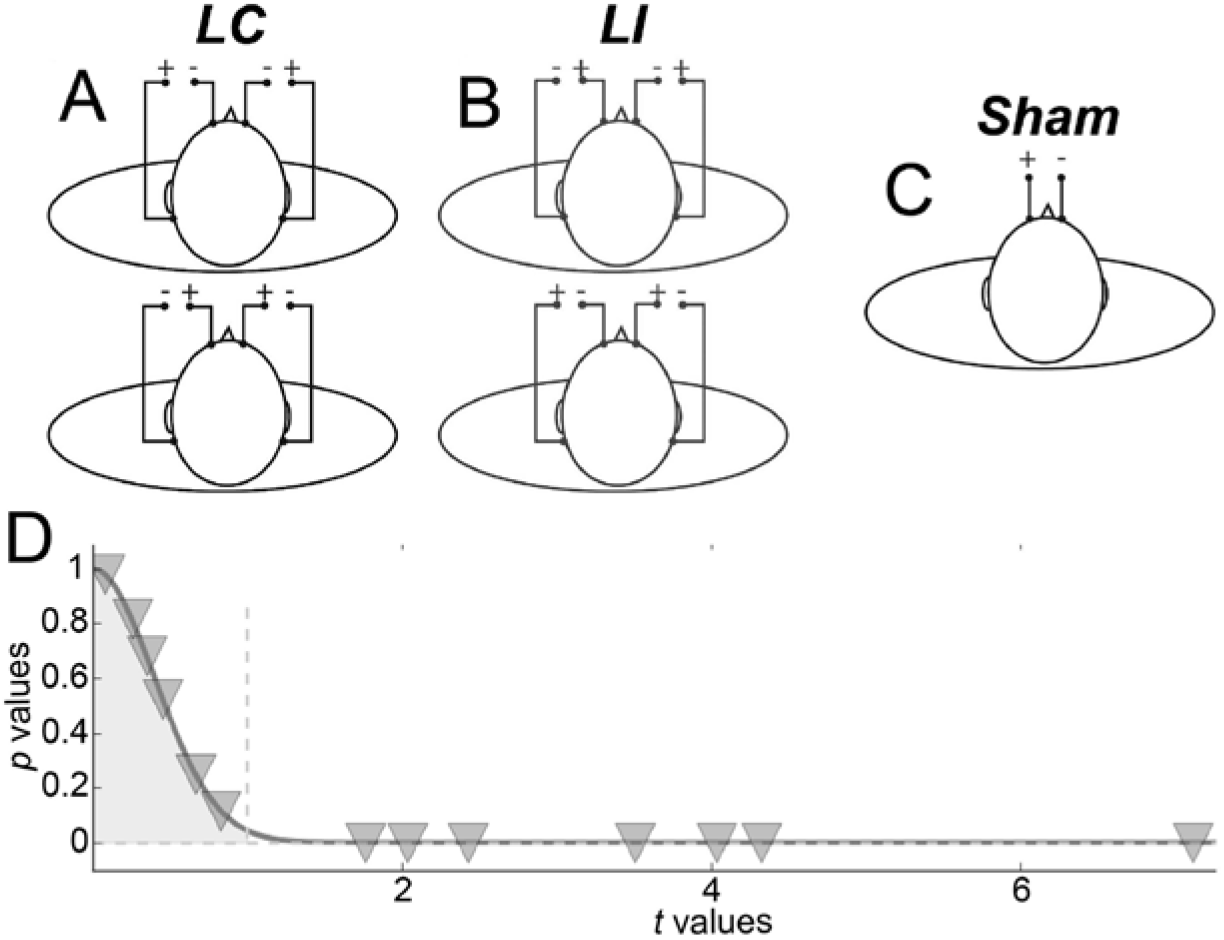
Galvanic vestibular stimulation (GVS) procedure (A) The 2 antero-posterior configurations. (B) The 2 lateral configurations. (C) Sham configuration. (D) Representation of behavioural results. Inverted triangles give the *p* and *t* values of the permutation tests for each subject. The vertical dashed line corresponds to the 95% confidence interval.

The stimulators were localized outside the scanner room and were connected to screen cables through a waveguide. Four GVS configurations were used. Two bilateral monopolar configurations with anodes over the mastoids (figure 1A, top) or the forehead (figure 1A, bottom) respectively permitted to elicit vestibular activations consistent with forward or backward motion of the body (antero-posterior GVS). Two bilateral bipolar conditions with anode right / cathode left (figure 1B, top) or cathode right / anode left (figure 1B, bottom) permitted to evoke activations consistent with leftward or rightward motion (lateral GVS). The amplitude of the postural reaction is known to vary with GVS intensity, until reaching a plateau (Séverac Cauquil et al., 2013). For that reason, on top of obvious avoidance of tactile or even painful stimulation, we chose to use low, 1mA, GVS intensity. Therefore, because the subjects were not always aware of the stimulation, a beep sound informed them every time a stimulation was delivered. As a baseline, we used a no-GVS condition that started with a beep but without any stimulation. Data were collected using an event-related design within which runs lasted 5 minutes (300s) and comprised 40 events (8 for each of our 5 conditions). The time interval between two condition onsets was fixed to 7.5s. Subjects were instructed to perform a forced-choice task using a 4-button box. After each beep, they had to press either the left or the right button to report whether they had experienced a sensation of self-motion along the lateral axis (L) or either the up or the down button in case of antero-posterior, AP, self-motion. The statistical significance of these L vs AP responses was evaluated for each subject through permutations tests (Figure 1D). For these permutation tests, we computed 10.000 synthetic means by randomly subsampling 27 trials from the 54 of the sham condition. We generated representative distributions of these mean values. A z-score and its corresponding p-value were then obtained by dividing the observed mean for the subjects in the stimulation trials by the standard deviation of the Gaussian distribution generated by the permutation tests (and always centred on ~1).

To control that our fMRI results were caused by vestibular activations rather than by somatosensory effects induced by the galvanic stimulation, we also designed a sham condition during which the same stimulation (i.e., a 2s square pulse of 1mA) was only delivered between the two frontal electrodes (figure 1C). Responses to these stimulations were recorded during a separated run of 5 minutes that comprised 40 events whose onsets were separated by 7.5s. The GVS and sham runs were conducted in total darkness on subjects instructed to keep their eyes closed during the whole recording to avoid any visual stimulation. We discuss the possible implications of eye movements on our results in the *‘Control for vergence*’ section.

#### Localizers for areas responding to egomotion-compatible optic flow

In this study, our main analyses were performed within functionally defined ROIs that preferentially respond to egomotion compatible optic flow. This ROI-based approach enables us to directly compare ROI data across subjects. It also gets rid of the multiple comparisons problem because statistics are only performed within the predefined ROIs (see e.g. Poldrack, 2007). In order to localize the cortical areas that respond to egomotion-compatible optic flow, we used the stimuli described in previous studies, see e.g. (Wall and Smith, 2008; Cardin and Smith, 2010). It consisted of 500 moving white dots displayed at 60Hz on a black background and arranged in an egomotion-consistent (EC) or egomotion-inconsistent (EI) optic flow pattern. In the EC condition, the optic flow pattern had both expansion/contraction and rotation components that varied over time, consistent with self-motion on a varying spiral trajectory (Morrone et al., 2000). The EI stimulus consisted of a 3 × 3 array of 9 identical panels, each containing a smaller version of the EC stimulus. Although the individual panels contained optic flow, the overall pattern was not consistent with egomotion because flow induced by observer motion can have only one centre of motion. Stimuli were presented using a block-design. Runs consisted of 224s (3min, 44s) divided into 7 identical cycles of 32s. In half of the runs, a cycle started with a baseline of 10s where only the fixation point was present. It was followed by 6s of the EC condition, then by another 10s of blank and finally by 6s of the EI condition. In the other half of the runs, the EC and EI conditions were inverted within a cycle (i.e. a cycle had 10s of blank, 6s of the EI condition, 10s of blank and finally 6s of the EC condition). During the recordings, subjects were instructed to passively keep their eye on the central fixation point. They, however, all reported that the EC conditions elicited a strong percept of egomotion.

The localizers for the ROI responding to egomotion-compatible optic flow were presented via an LCD projector, back projected onto a screen positioned at the end of the scanner bore, and viewed through a mirror mounted on the head coil. The viewing distance was 130cm. It led to squared stimuli of 16°x16°.

#### Data acquisition

All the data were collected on a 3T scanner (Philips Achieva), using a standard 32 channels head coil. The functional data were acquired using (T2*-weighted) echoplanar imaging (EPI). The data for the main experiment (GVS) were collected during a first session. The data for the functional localizers were collected during a second session.

For the GVS experiment, we used the following prescription that is quite generic for whole-brain recordings: time repetition (TR) = 2.5 s, time echo (TE) = 30 ms, voxel size 3×3×3 mm, no gap thickness, flip angle (FA): 77°, SENSE factor: 2.8. Each run comprised 120 volumes of 41 transversally oriented slices that covered the whole brain. In total, we collected 10 runs (8 runs with the 4 main conditions and the baseline and two additional runs with the sham stimulations, see the *‘Galvanic stimulation*’ section above). The total duration of the recordings was about 45 minutes.

For the functional localizers, because the cortical regions that selectively respond to egomotion consistent optic flow are now well established in the occipital and parietal regions (see e.g. Cardin & Smith, 2010 or Smith et al., 2012), we used a prescription specifically designed to optimize the resolution of BOLD recordings in these particular regions: TR: 2 s; TE: 30 ms; field of view (FOV): 210 mm; voxel size 2×2×2mm; no gap thickness, SENSE factor: 2.5. A run comprised 96 volumes of 33 slices that covered occipital and parietal cortices. We recorded 4 runs in total (2 for each condition).

Both the two sessions of recordings also included the acquisition of a high-resolution anatomic image using a T1-weigthed magnetization-prepared rapid gradient-echo (MPRAGE) sequence (160 slices; TR: 2300 ms; TE: 3.93 ms; FA: 12°; FOV: 256 mm; voxel size 1×1×1 mm). These anatomical images were first co-registered and then averaged together to be used as a reference to which the functional images from all the experiments were aligned.

### Data Analyses

#### Pre-processing

All the fMRI data were analysed using the Brain Voyager QX software (v2.8, Brain Innovation) and Matlab. Pre-processing included slice scan time correction, 3D motion correction using trilinear/sinc interpolation, and high-pass filtering (0.01 Hz). For each individual subject, functional data were co-registered on the anatomy. Functional and anatomical data were brought into ACPC space using cubic spline interpolation and then transformed into standard Talairach (TAL) space (Talairach and Tournoux, 1988).

#### Region of Interest (ROI) definition

For each subject who performed the localizers experiment (n = 11), we determined the areas responding to egomotion compatible optic flow (V6, VIP, CSv, hMT+ and PIC) using the contrast between egomotion-consistent (EC) *vs* inconsistent (IC) optic flow conditions (see the ‘*Localizers for areas responding to egomotion-compatible optic flow*’ section above). Except for area V6 for which we used the adaptive statistical threshold procedure proposed in Cardin et al. (2012b)(see below), our functional areas were defined using a threshold of p < 0.001 (uncorrected).

V6 seed was determined as the most significant voxel within the parieto-occipital sulcus (POS) for the EC *vs* EI contrast. We then grew a V6 cluster around this seed by reducing the threshold until the point in which the cluster started to expand outside the POS (Cardin et al., 2012b). We defined V6 at this threshold. This approach led to a successful identification of area V6 in 10 out of our 11 subjects who underwent the localizers. Because our V6 ROI was not defined from wide-field retinotopic mapping (see e.g. Pitzalis et al., 2006), we cannot be certain about the exact limit of this ROI. We therefore propose a control analysis to determine if this uncertainty impacts our results (see the ‘Results’ section).

Using the same contrast between egomotion-consistent (EC) *vs* inconsistent (EI) optic flow, we also defined the ventral intraparietal (VIP) area. This was the cortical region in the anterior part of the intraparietal sulcus and close to the intersection with the post-central sulcus that was significantly more activated during the EC condition. This location matches with the one reported in the original study of Bremmer et al., (2002), and is consistent with the definition of VIP described in Smith et al., (2012). Using this definition, we were able to define VIP bilaterally in 7 of our subjects. For another 3 subjects, we localized VIP in one hemisphere but not in the other. Then, for each subject, the data corresponding to an ROI that was found bilaterally were averaged across hemispheres.

With the same approach, we defined area CSv in all our subjects and the human middle temporal complex hMT+ in 10 of our 11 subjects. This region was localized within the ascending branch of the inferior temporal sulcus (ITS) (see Kolster et al., 2010) and includes MT, MST and possibly other few motion regions like the putative fundus of the superior temporal area (pFST). Finally, our contrast also revealed a visually responsive region in the vicinity of parieto-insular cortex (PIC) in 9 of our subjects. This region corresponds to an area originally described by Sunaert et al., (1999), and that responds more strongly to the egomotion-consistent stimuli (see Billington and Smith, 2015). PIC was recently proposed as a putative homolog of macaque VPS (Frank et al., 2014).

The average TAL coordinates of these ROIs in all our subjects, provided in table 1, fit very well with those reported in previous studies. Some coordinates from these previous studies, indicated by an asterisk in the table, were transformed from MNI to TAL to allow direct comparison with ours. All the identifications of ROIs in this study where performed using WFU Pickatlas, version 3.05 (ANSIR Laboratory, WFU School of Medicine, NC-USA) (Lancaster et al., 2000; Maldjian et al., 2003). Figure 2-A shows the results of the contrast between egomotion-consistent (EC) versus inconsistent (EI) optic flow and the resulting ROIs in one typical participant.

**Table 1:** ROI comparison with previous studies, *originally published in MNI coordinates and transformed to Talairach using WFU pick atlas (Lancaster et al., 2000; Maldjian et al., 2003)

| Area | Mean Coordinate (TAL) | Reference (TAL) | Study |
| --- | --- | --- | --- |
| Left V6 | -13 -81 27 | -11 -79 30 | (Cardin and Smith, 2010) |
| Right V6 | 15 -76 30 | 14 -77 30 |  |
| Left VIP | -45 -41 38 | -40 -40 42 | (Bremmer et al., 2001) |
| Right VIP | 41 -48 40 | 38 -44 46 |  |
| Left CSV | -10 -25 40 | -10 -25 38 | (Wall and Smith, 2008) |
| Right CSV | 11 -26 41 | 10 -26 41 |  |
| Left MT+ | -44 -62 5 | -39 -62 5 | (Cardin and Smith, 2010) |
| Right MT+ | 43 -62 3 | 39 -60 -1 |  |
| Left PIC | -40 -21 21 | $\pm 44$ -28 23 | (Sunaert et al., 1999) |
| Right PIC | 41 -22 19 |  |  |

**Figure 2:**
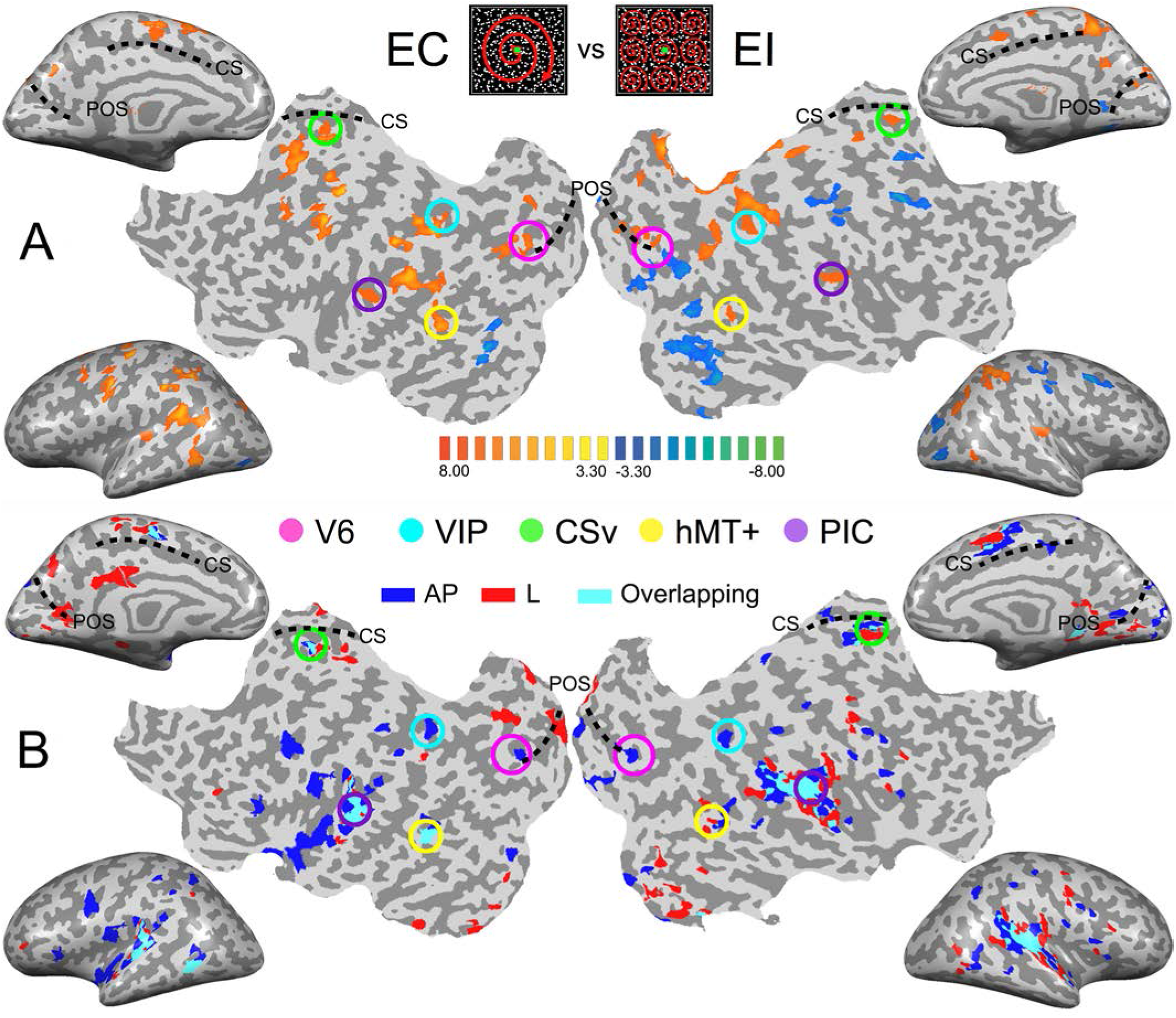
Images from the brain of one participant showing the key results. (A) Contrast between the cortical responses recorded during the egomotion consistent (EC) versus inconsistent (EI) optic flow conditions. Data are shown on inflated cortical surfaces and flat maps for the left and right hemispheres (p<0.001, uncorrected). The positions of the parieto-occipital sulcus (POS) and Cingulate sulcus (CS) are provided as anatomical landmarks. Colored circles outline the five ROIs in this subject. (B) Egomotion-consistent (EC) and egomotion-inconsistent (EI) optic flow patterns used as localizers stimuli (C) Responses during the galvanic stimulation (GVS) experiment for the same participant. Voxels whose activations were stronger during GVS conditions than during baseline were colored in blue (antero-posterior, AP), red (lateral, lat) and cyan (both) before being superimposed transparently on the inflated brain and flat maps.

#### General linear model (GLM)

All our analyses of the functional data used a general linear model (GLM). The data from each participant were analysed separately. Time series were processed by fitting a regressor formed by convolving the event time course with a standard hemodynamic response function. As GVS induced micro-movements of the participant’s head could potentially bias our results, six regressors taken from the head motion correction were also included as regressors of no interest. The responses to our first and second conditions (i.e. the stimulations that elicited a vestibular activation consistent with a forward or a backward motion) were modelled together to form the responses to antero-posterior (AP) stimulations. The responses to our third and fourth conditions (i.e., the stimulations consistent with a leftward or a rightward motion) were also modelled together to form the responses to lateral (lat) stimulations. Finally, the beta values obtained for these two conditions and for the sham stimulation were corrected by subtracting the beta values obtained during the baseline condition. Then, we looked at the results at the individual level. Our analysis focuses on ROIs that were specifically involved in the processing of EC optic flow to check whether they also had specific responses to AP or Lat galvanic stimulations. However, we also completed this approach with a preliminary whole brain analysis that was performed at the individual level. In this case, activations were first displayed as an overlay of a segmented and inflated or flattened representation of each hemisphere based on the average anatomical scan of each subject. Activation maps were thresholded at p<0.001 (uncorrected). The aim of this initial whole brain analysis was to obtain a general overview of our data and thereby to avoid pinhole conclusions (see Hupé, 2015).

#### Connectivity analyses

To characterize functional connectivity between our ROIs during our two conditions, we performed a psychophysiological interaction (PPI) analysis (Friston et al., 1997). This analysis aims at characterizing task or context specific changes in the relationship between brain areas (see e.g. O’Reilly et al., 2012 for a review). In our specific case, it permitted to establish those cortical areas that are specifically more connected during the AP and Lat stimulations. PPI can be obtained with a general linear model (GLM) that contains three regressors: the psychological variable (in our case antero-posterior/lateral, coded as +1/-1), the physiological variable (the time-course of a seed region) and the PPI regressor (psychological × physiological regressor). Before computing the interaction term, the psychological and physiological time courses were both expressed in terms of the underlying neural activity. To do so, we first estimated the hemodynamic response function and then used it to deconvolve the activity recorded from the seed ROI (Gitelman et al., 2003). The two time-courses (psychological and physiological) were also included in the GLM as co-variates of no interest. This means that variance explained by the interaction term is only that over and above what is explained by the main effects of task and physiological correlation. We constructed one GLM for each of our ROIs. The seed time-course associated with an ROI first corresponded to the average response of the ROI across its voxels. It was then mean-corrected and z-transformed. The psychological variables were the AP GVS condition *vs* baseline on the one hand and the Lat condition *vs* baseline on the other hand. The PPI predictor of a given seed region was then tested in each of the remaining network nodes in a multisubject RFX GLM (points 3-5, covering the peak of the BOLD response).

To focus our analysis on the connections within our functionally defined ROIs, we performed a multiregional PPI approach (Cocchi et al., 2014, Schindler and Bartels, 2016). Multiregional PPI is a simple generalization of the PPI approach; it permitted to characterize connectivity between each pair of our functionally defined ROIs (10 pairs in total) rather than between a single-seed region and all the other brain voxels. This analysis was performed at the single subject level. We, therefore, performed 9 analyses corresponding to the 9 subjects for whom we were able to identify all the ROIs. The PPI predictor of a given ROI was then tested in each of the remaining network nodes in a *ROI-paired* multisubject RFX GLM (points 3-5, covering the peak of the BOLD response).

## Results

### Behavioural results

We analysed the behavioural responses collected for each subject. Although it is well-established that the galvanic stimulation configuration (monopolar *vs* bipolar) has a significant effect on the perceived direction of self-motion in standing and lying subjects (Fitzpatrick et al., 2002, Fitzpatrick and Day, 2004; Lepecq et al, 2006; St George et al, 2011; Ferrè et al., 2013; Aoyama et al., 2015), we did not necessarily expect to elicit clear sensations in our experiment because of the short stimulation duration and low intensity used in our design (see the *‘Galvanic stimulation’* section). At the group-level, we did not find significant differences between the behavioural responses to our AP and Lat conditions (*paired t-test t*_13_ = 0.16, *p*= 0.87). Nonetheless, at the individual level, we found that 7 of the 13 subjects were able to significantly discriminate between the AP and Lat conditions (Figure 1D).

### Whole Brain analysis

As an initial step, we computed for each subject the activation maps during the galvanic stimulation (GVS) conditions using a whole-brain analysis. This enables us to obtain an overview of the data at the individual level and also to compare the maps across subjects. However, bear in mind that our analysis (at both the individual and the group level) is performed within our functionally defined ROI (see the ‘ *General Linear Model* (*GLM*)’ section of the Materials and Methods and the next section). For this initial step, we contrasted both the antero-posterior (AP) and lateral (lat) GVS conditions against the baseline. These contrasts for one typical participant are shown in figure 2-B (p<0.001, uncorrected). For a direct comparison between visual and vestibular responses, figure 2-A also shows response to optic flow in the same participant.

The responses to lateral GVS (in red) are in good agreement with previous imaging studies that used similar GVS conditions (Bucher et al., 1998; Lobel et al., 1998; Bense et al., 2001). Activity was seen in the parieto-insular vestibular cortex (PIVC) and in putative vestibular areas 2v and 3aNv. Consistently with the previous work of Smith et al. (2012) (see their figure 2), we also found activations in visual areas such as the hMT+ complex or CSv, a portion of the cingulate sulcus that is highly activated during the presentation of egomotion-compatible optic flow-see e.g. Wall & Smith (2008), or Smith et al. (2012). Responses in the other participants were very consistent with those observed here.

Across subjects and hemispheres, responses to antero-posterior GVS stimulation were generally similar to those observed during lateral GVS stimulation. However, the former condition led to stronger responses in several cortical regions. One is located within the posterior part of the parieto-occipital sulcus. Another lies within intra-parietal sulcus, close to its intersection with post-central sulcus. These two regions overlap with our functionally defined ROIs V6 and VIP (see figure 2-A, the pink and cyan circles). Outside our visual ROIs, we did not find any region that was consistently (i.e., across subjects and hemispheres) more activated by one of our two GVS conditions.

### Regions of Interest analysis

We ran a ROI-based analysis to enable the comparisons between the responses from our different subjects (see the ‘Materials & Methods’ section). Within all the ROIs (in both hemispheres) of our subjects, we computed the beta values corresponding to the AP GVS, Lat GVS and the sham condition. These beta values were then corrected by subtracting the beta values of the baseline condition (see the *‘General Linear Model (GLM)’* section). Figure 3 shows the results in all our ROIs (i.e., V6, VIP, CSv, hMT+ and PIC).

**Figure 3:**
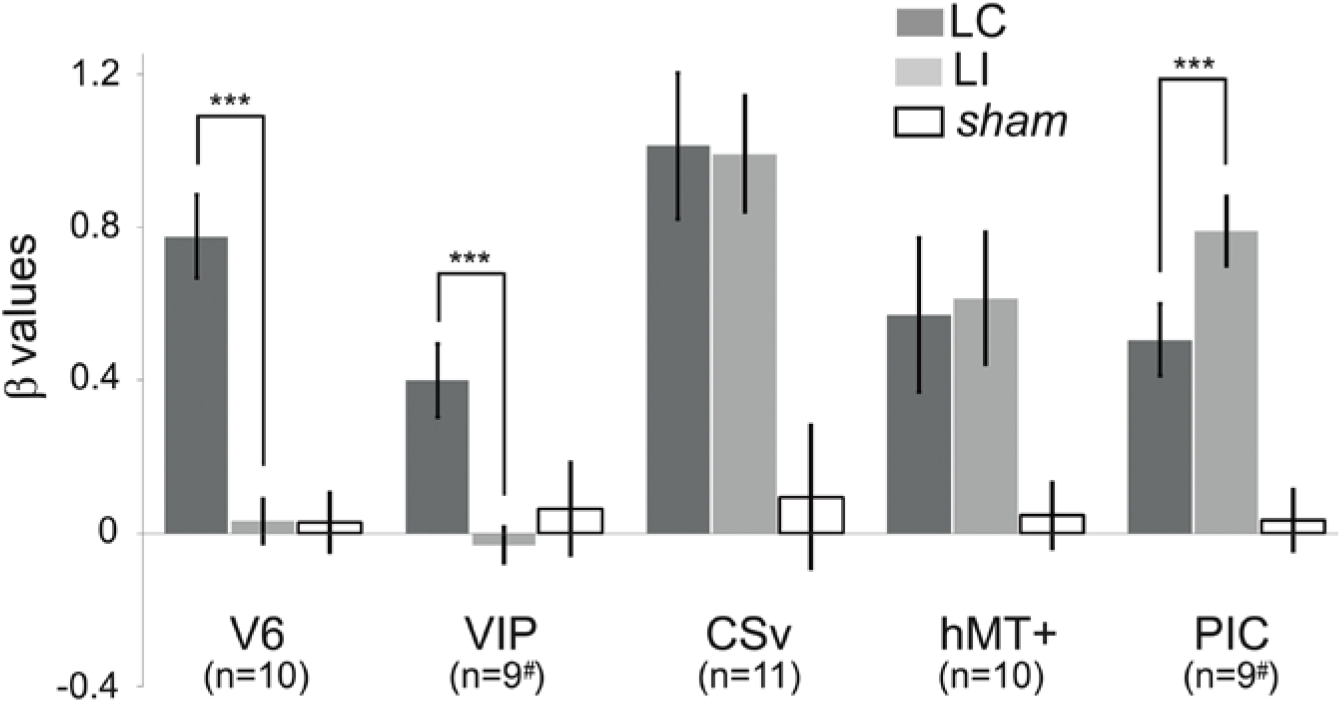
Average beta values obtained in our different ROIs during the antero-posterior (dark grey) and lateral (light grey) GVS conditions. Values corresponding to the sham condition are provided in white. The error bars give the standard errors. The # symbols are here to remind that in some subjects, the VIP and PIC ROIs were only defined in one hemisphere (see details in the text). We report here the significant differences between AP and Lat conditions (post-hoc t-test, ***: p<0.001, *: p<0.05). Results of the other statistical comparisons are reported in the main document.

Responses in V6 were strongly dependent of the condition (rmANOVA, Greenhouse-Geisser corrected: F(2)=32.02, p<0.001, η^2^=2.81) (figure 3, table 2). Post-hoc *t-tests* confirmed that the beta values were significantly higher for the AP GVS condition (t(9)=6.99, <0.0001 when compared to the Lat condition and t(9)=7.72, p<0.0001 when compared to the *sham* condition). We did not find any significant differences between the Lat condition and the *sham* condition (t(9)=0.046, p=0.964). Responses were also strongly influenced by condition in area VIP (rmANOVA, Greenhouse-Geisser corrected: F(2)=5.52, p=0.016, η^2^=0.512). In this ROI as well, post-hoc *t-tests* showed that responses in the AP GVS condition were stronger that in the Lat condition (t(9)=4.252, p<0.05). We did not find any significant differences between the Lat condition and the *sham* condition (t(9)=0.667, p=0.521). Therefore, both areas V6 and VIP had specific responses during the antero-posterior GVS.

**Table 2.** Paired t-test results, contrasting beta-values obtained in AP, Lat, and sham condition, for our 5 Regions Of Interest, by taking all (100%) voxels, and the 80% and 60% closest to the central coordinates.

| 100% |  |  |  |
| --- | --- | --- | --- |
| AREA | AP-LAT | AP-SHAM | LAT-SHAM |
| V6 | t(9)=6.99, p=0.000064 | t(9)=7.72, p=0.000029 | t(9)=0.046, p=0.964 |
| VIP | t(9)=4.252, p=0.021 | t(9)=2.625, p=0.029 | t(9)=0.667, p=0.521 |
| CSV | t(10)=1.03, p=0.91 | t(10)=4.22, p=0.002 | t(10)=4.24, p=0.02 |
| hMT+ | t(9)=-0.145, p=0.887 | t(9)=2.83, p=0.019 | t(9)=3.91, p=0.003 |
| PIC | t(9)=-2.41, p=0.0269 | t(9)=4.193, p=0.002 | t(9)=6.86, p=0.000081 |
| 80% |  |  |  |
| AREA | AP-LAT | AP-SHAM | LAT-SHAM |
| V6 | t(9)=4.07, p=0.003 | t(9)=2.96, p=0.016 | t(9)=0.738, p=0.479 |
| VIP | t(9)=2.503, p=0.034 | t(9)=4.391, p=0.002 | t(9)=2.705, p=0.026 |
| CSV | t(10)=0.386, p=0.708 | t(10)=3.416, p=0.007 | t(10)=4.399, p=0.001 |
| hMT+ | t(8)=0.247, p=0.811 | t(8)=4.04, p=0.003 | t(8)=5.012, p=0.001 |
| PIC | t(9)=2.705, p=0.024 | t(9)=4.256, p=0.002 | t(9)=6.36, p=0.000131 |
| 60% |  |  |  |
| AREA | AP-LAT | AP-SHAM | LAT-SHAM |
| V6 | t(9)=2.822, p=0.02 | t(9)=2.545, p=0.027 | t(9)=1.101, p=0.3 |
| VIP | t(9)=2.872, p=0.018 | t(9)=4.923, p=0.000821 | t(9)=2.339, p=0.044 |
| CSV | t(10)=0.555, p=0.592 | t(10)=2.301, p=0.04 | t(10)=3.126, p=0.011 |
| hMT+ | t(7)=0.073, p=0.944 | t(7)=3.346, p=0.012 | t(7)=3.394, p=0.011 |
| PIC | t(9)=0.187, p=0.856 | t(9)=4.574, p=0.001 | t(9)=6.254, p=0.000149 |

Responses in area CSv and hMT+ were both strongly modulated by condition (rmANOVA, Greenhouse-Geisser corrected: F(2)=3.253, p<0.001, η^2^=3.59 in CSv and F(2)=4.522, p<0.05, η^2^=1.429 in hMT+). In these ROIs, both the AP (t(10)=4.22, p<0.01 in CSv and t(9)=2.83, p=0.05in hMT+) and Lat (t(10)=4.24, p<0.01 in CSv and t(9)=3.91, p=0.05 in hMT+) GVS conditions had stronger responses than the sham condition. This time, we did not find any significant difference between the two GVS conditions (t(10)=1.03, p=0.91 in CSv and t(9)=-0.145, p=0.887 in hMT+).

Finally, responses in PIC were also dependent of the condition (rmANOVA, Greenhouse-Geisser corrected: F(2)=55.55, p<0.001, η^2^=1.579). In this case, post-hoc t-tests confirmed that responses were stronger during the Lat GVS condition than during the AP GVS condition (t(9)=-2.41, p<0.05). Responses during the AP and Lat conditions were stronger than during the *sham* condition (t(9)=4.193, p<0.01 and t(9)=6.86, p<0.001 respectively). Among all our ROIs, CSv had the strongest responses to GVS conditions. This result is consistent with those reported in Smith et al., (Smith et al., 2012). Overall, our results demonstrate that the activations elicited by Lat GVS in areas V6, VIP, CSv, hMT+ and PIC are reliable across different stimulation parameters (a 1Hz sinewave alternating between +/-3mA in Smith et al., 2012 versus a 1mA step in the present study).

We observed in the *‘Region of Interest (ROI) definition’* section that our procedure to define area V6, which did not include wide-field retinotopic mapping, cannot guarantee that this ROI does not include small portions from adjacent areas in some subjects. To make sure that the effects reported here reflect properties of area V6, we performed a control analysis where we reproduced our statistics on subsamples of voxels within this ROI. We first computed the Euclidean distances between all the voxels within the ROI and the ROI center. We then defined two smaller ROIs that grouped either the 80 or the 60% of voxels that were the closest to the central coordinates. These smaller ROIs have less chance to contain voxels that do not belong to V6.

The results of this analysis are shown in figure 4 and table 2 reports all the paired t-test contrasts between the GVS conditions, for our 5 ROIs.

**Figure 4:**
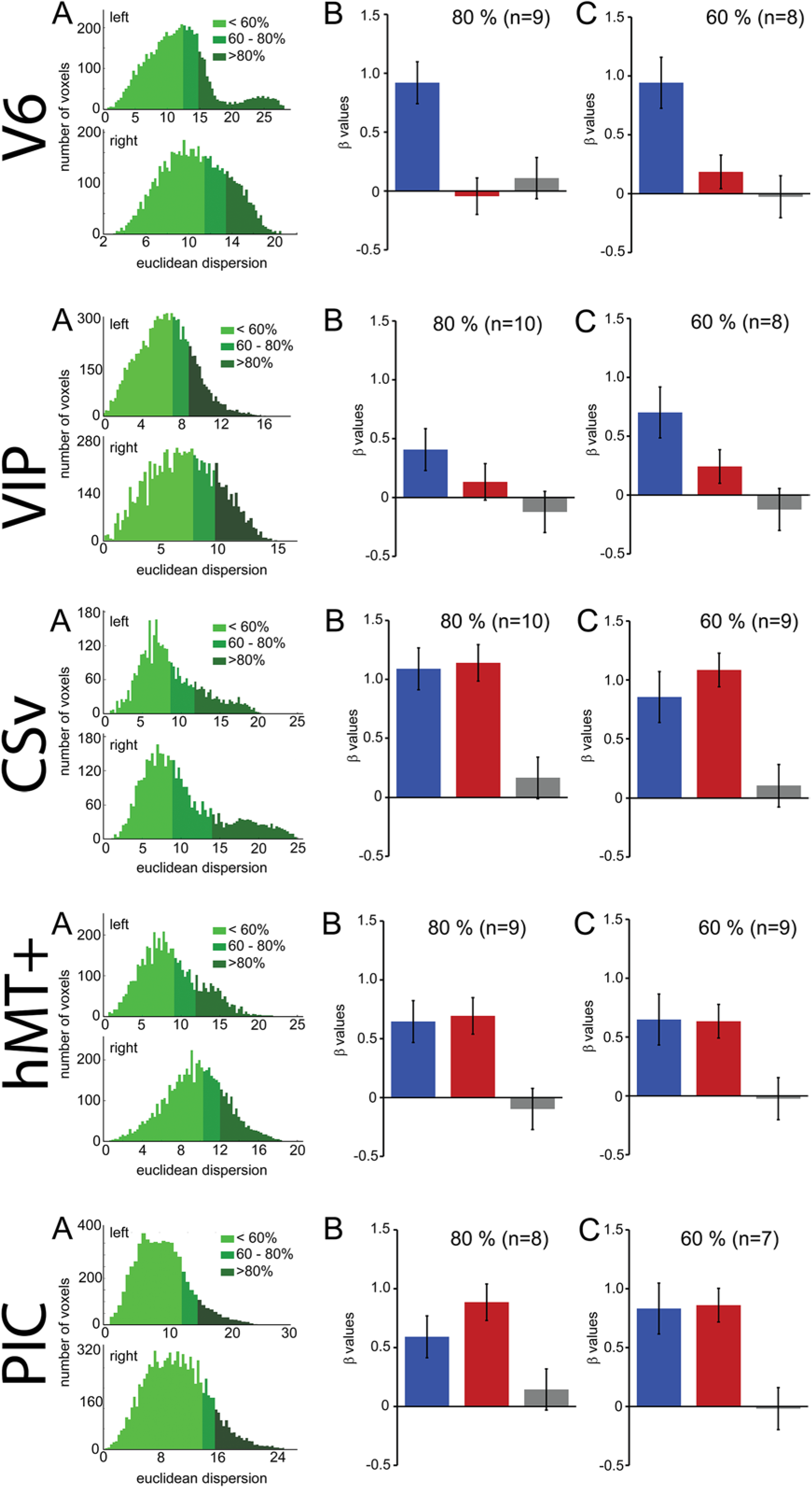
Control to characterize the influence of the ROI spatial extents on the results. These analyses were performed for V6, CSv, hMT+, VIP and PIC (from top to bottom). (A) Histograms showing the number of voxels within an ROI as a function of the Euclidean distance to the ROI centroid. The green colours give the repartition of the 60%, 80% and 100% closest voxels. (B) Bar graphs of the beta values for the two GVS conditions (AP, in blue and Lat, in red) and the sham stimulation. The analysis only included the 80% closest voxels to the centroid. (C) Idem for the 60% closest voxels.

We can observe that our main results (figure 4, table 2) remained unchanged with this analysis. It demonstrates that the preference for AP GVS stimulation that we found in V6 is robust to variation in the spatial extent used to define this area and, therefore, not driven by activity within adjacent functional areas. We used the same approach to double-check our results in the VIP, CSv, hMT+ and PIC ROIs. Indeed, these areas were defined from thresholded contrast maps (p < 0.001, uncorrected, see the Materials and Methods), which is always subject to uncertainty, see e.g. Eickhoff et al. (2009). This control analysis confirmed our results in areas CSv, hMT+ and VIP. In particular, for area VIP, it demonstrated that the preference for AP GVS did not depend on the spatial extent used to define this ROI. Interestingly, our control confirmed that responses in PIC during Lat GVS were stronger than during AP GVS only for the 80% of voxels but not for the 60% (t(9)=2.705, p=0.024 and t(9)=0.187, p=0.856 respectively). This result should therefore be taken with care and will probably necessitate further investigations.

### Connectivity analysis

The differential activation within our visual ROIs during antero-posterior, (AP) and lateral (lat) conditions supports the hypothesis that antero-posterior and lateral vestibular signals are processed by distinct cortical networks. Nevertheless, it does not provide any information regarding interactions between these areas and the structure of these networks. In order to identify the connectivity pattern between our functionally defined ROIs, we ran a *multiregional psychophysiological interaction* (PPI) analysis (see the *‘Connectivity analysis’* section). For this type of analysis, it is mandatory that all ROIs are defined in each subject. It was, therefore, only performed on the 9 subjects for whom all the ROIs were defined. If V6 and VIP are more activated during the AP compatible condition, one could expect that connectivity between each of these ROIs and the others are more pronounced during this condition. Figure 5-A shows connections between our ROIs that are significantly more correlated during the Lat condition than during baseline. Figure 5-B shows these correlations for the AP condition.

**Figure 5:**
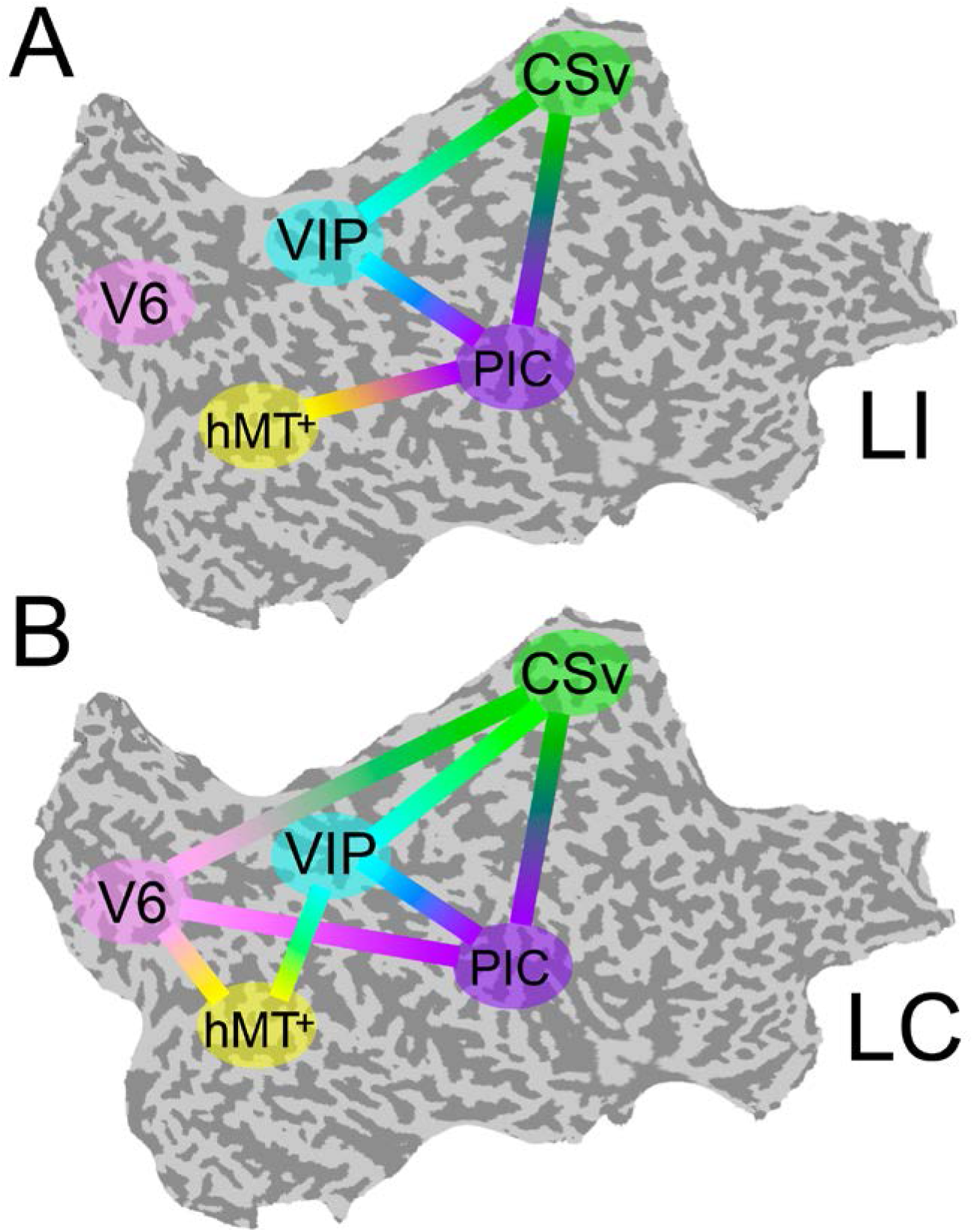
PPI functional connectivity analysis. Multiregional PPI was run across the 5 ROIs identified in the localizer protocol. Results are shown for the Lat (A) and AP (B) GVS conditions. A solid line between 2 ROIs corresponds to a connection that is significantly stronger during this condition (P<0.001) than during baseline.

During Lat GVS, areas PIC, hMT+, CSv and VIP were more connected together than during baseline (5-A). This was particularly true for area PIC. The connectivity pattern between V6 and the other ROIs remained at the baseline level for this condition. At the opposite, during AP GVS, V6 was significantly more connected to CSv, hMT+ and PIC (5-B) than during baseline (P<0.001). Area VIP was also significantly more connected to hMT+ during this condition (P<0.001), whereas it was not the case during the Lat condition.

In order to determine if cortical activations were different in the subjects who detected the GVS direction over chance, we looked for correlation between behaviour and brain activity. We did not find any significant relationship between our fMRI measurements and our subject’s perceptual reports. This was true for both the beta values and the connectivity strengths. Given our short stimulation duration and the low intensity used in our design, it is possible that the elicited percept was not strong enough to establish such correlations.

### Control for vergence

Our results showed that areas V6 and VIP are only activated during antero-posterior GVS. One possibility is that, even if our participants had their eyes closed, this condition triggered convergence or divergence eye movement and these movements affected the activity in V6 and/or VIP. For example, a study by Quinlan and Culham (2007) showed that responses in the dorsal parieto-occipital sulcus (dPOS, a brain portion that includes V6) were modulated by the vergence angle. To control that GVS (and specifically the stimulations associated with a backward or forward motion of the body) did not trigger convergence and/or divergence movement of the two eyes, we performed a control experiment outside of the scanner. For this control, subjects had their eyes opened and binocularly viewed Nonius lines (see Cottereau et al., 2011) through anaglyph goggles with red/green filter on the left/right eye. The green line was displayed in the upper part of the visual field and was only seen by the right eye (through the green filter). The red line was displayed in the bottom part of the visual field and was only seen by the left eye (through the red filter). The two lines were vertically aligned with a visible fixation point at the center of the screen. When binocularly viewed through the anaglyph goggles, this configuration appeared as two white lines vertically aligned with a white dot on the centre of the screen. The subject task was to fixate the point during blocks of GVS that were identical to those used in our main experiment (see the ‘Materials and methods’ section). After each stimulation, subjects had to report if they perceived the two lines as ‘aligned’ during GVS (upper arrow of the keyboard) or if the upper line moved to the left (left arrow) or to the right (right arrow) relatively to the bottom line. These last two cases, respectively, correspond to convergence and divergence eye movements. The sensitivity to Nonius misalignment is typically below 2 arcmin (McKee and Levi, 1987), which is more accurate than what can be obtained from a binocular eye tracker. 5 subjects who participated in the galvanic stimulation experiment performed 20 trials of each condition. Their perceptual reports are provided in figure 6.

**Figure 6:**
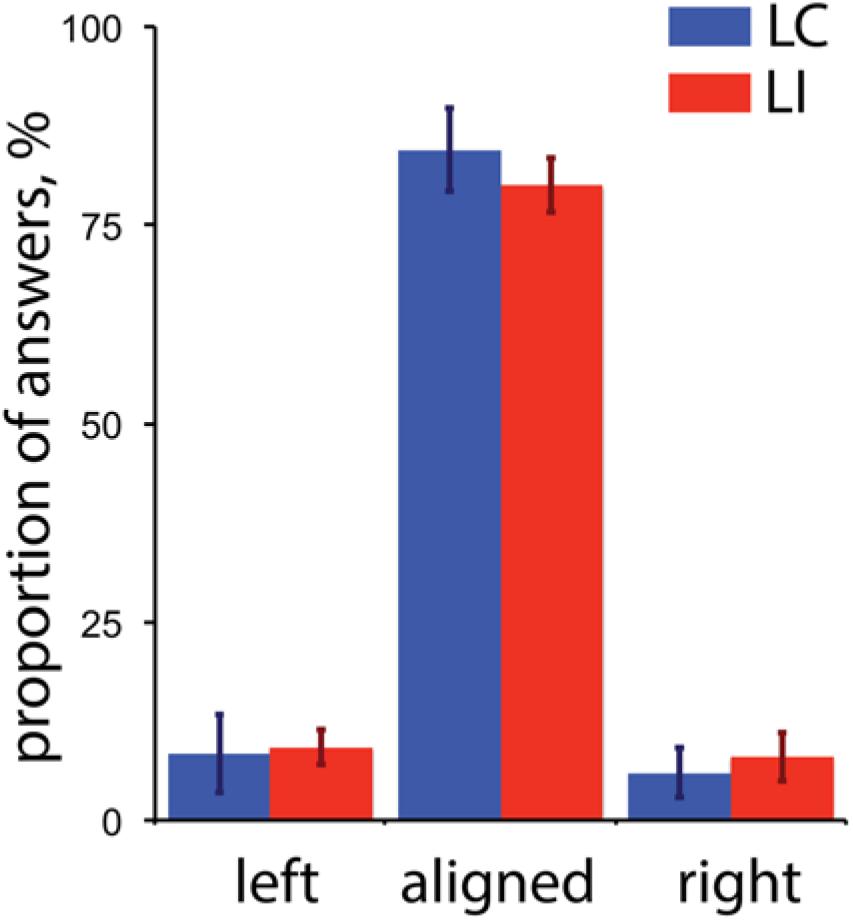
Perceptual reports during the control experiment for vergence (n = 5). The proportion of ‘left’, ‘aligned’ and ‘right’ answers are provided for both the antero-posterior (AP, blue) and lateral (Lat, red) conditions. The ‘left’ and ‘right’ reports respectively correspond to convergence and divergence eye movements (see details in the text). The error bars give the standard errors.

These results demonstrate that: 1) GVS had a minor impact on binocular eye movements, and 2) the small proportions of reported convergence/divergence movements were not statistically different between our AP and Lat GVS conditions. We conclude that the activations elicited by our AP condition in V6 were not caused by eye movements.

Even though our subjects had their eyes closed during the fMRI recordings, we cannot exclude that their lateral eye movements were different between our two conditions. If it is a limitation of our study, we are confident that our results are not contaminated by lateral eye movements. Indeed, our analyses were performed in functional ROIs that are not specifically known to respond to lateral eye movements. In addition, our whole brain analysis did not reveal any significant activation in regions whose responses are actually modulated by lateral eye movements like for example the frontal eye field.

### Discussion

The aim of this study was to characterize the cortical networks that are activated during antero-posterior (AP) and lateral (Lat) GVS using fMRI measurements. A previous neuroimaging study employed the usual binaural bipolar mode, where the anode is placed on one mastoid process and the cathode on the other to identify visual cortical areas that receive vestibular inputs (Smith et al., 2012). In the present study, we applied lateral GVS using an opposite double monaural configuration (see figure 1B) that was found to induce equivalent postural response than binaural bipolar (Séverac cauquil et al., 2000). Recent electrophysiology studies using GVS on macaque monkeys showed that anodal and cathodal have opposite effect on vestibular afferents discharge of both otolith and semicircular canals (Kwan et al., 2017). This corroborates the assumption that the orientation of the response to GVS is a function of the imbalance between right and left vestibular polarization (Séverac Cauquil et al., 2000). Here we replicated Smith’s results obtained from 3mA sinusoidal binaural bipolar stimulation using 1 mA step pulse in opposite double monaural GVS, validating the robustness of GVS approach. However, such lateral GVS configurations activate the parts of the vestibular apparatus that are sensitive to roll tilt, in the frontal plane (Day et al., 1997; Severac Cauquil et al., 2003; Fitzpatrick and Day, 2004). Therefore, this design prohibits the study of the consequences of an antero-posterior stimulation, although these signals are the most prominent during locomotion, which constitutes a major component of egomotion. We therefore used binaural monopolar GVS to investigate the cortical responses specific to antero-posterior mechanisms. This design, with electrodes of the same polarity placed over the mastoid processes, and of opposite polarity on the forehead orientates the galvanic-evoked vestibular input along the antero-posterior axis (Magnusson et al., 1990; Severac Cauquil et al., 1998, 2000; Aoyama et al., 2015), (see figure 1). This permitted us to distinguish the contribution of AP signals from the Lat ones provided by the usual, binaural bipolar mode. Our behavioural analysis supports those previously reported results. Even though our subjects were lying in the scanner, 7 over 13 were still able to discriminate over chance the stimulated direction of self-motion. Regarding the low intensity (1 mA vs 3 mA in Aoyama et al study (2015)) and short duration (2s vs 5s in Fitzpatrick and Day report (2002)) and taking into account the fact that here we submitted our subjects to a discrimination and not a detection task, we are entitled to consider we achieved to stimulate in two different directions our subjects’ vestibular apparatuses.

As a preliminary step, we performed for each individual subject a whole-brain analysis to get a general overview of our data. Across subject, our two GVS conditions led to strong fMRI activations within lateral sulcus, in the parieto-insular vestibular cortex (PIVC). This is in agreement with previous studies that found significant activations in the same region (Bucher et al., 1998; Lobel et al., 1998; Bense et al., 2001; Stephan et al., 2005). However, we were mostly interested in responses within functionally defined ROIs that are activated by egomotion-consistent optic flow: V6, VIP, CSv, hMT+ and PIC (see Cardin and Smith, 2010)(figure 2). Our aim was to better understand how these visual ROIs process vestibular inputs and hence their possible implication in multisensory integration during forward locomotion.

We found that area PIC was significantly activated during both our two GVS conditions (figure 3). This result is in agreement with a previous multisensory study (Frank et al., 2014). Actually, our connectivity analysis showed that PIC is the most connected area during galvanic stimulation (figure 5). This area possibly works as a hub where multisensory signals are integrated during egomotion. This strong selectivity to both visual and vestibular modalities supports the idea that PIC is the human homologue of macaque visual posterior sylvian area (VPS) (Chen et al., 2011). This is in total agreement with previous single-unit recordings and tracer studies in non-human primate (Guldin and Grusser, 1998). In macaque, this portion of cortex receives inputs from all the cortical areas of the vestibular system and also, even more relevant for our study, its neurons are sensitive to both somatosensory and visual signals, in particular to optokinetic stimulation from wide (i.e., > 30°) structures patterns (Grusser et al., 2010). Interestingly, it was found that if most vestibular neurons in PIC also respond to neck and visual stimuli, its processing of these signals does not appear to be done in a common reference system (Shinder and Newlands, 2014).

A major finding of this study is that area V6 is only activated during antero-posterior GVS (figure 3). Using lateral GVS, a previous study (Smith et al., 2012) did not find any activation in V6 and concluded that this area was probably not involved in visuo-vestibular integration. Our results are in agreement with the finding that V6 remains silent during lateral stimulation. However, the strong responses that we obtained during AP GVS show that V6 does receive vestibular input and has probably a specific role during locomotion. This hypothesis is strengthened by our PPI analyses that demonstrated that area V6 becomes significantly more connected to all our other ROIs during AP GVS (figure 5). A recent study showed that there is another visual region bordering V6: V6A (Pitzalis et al., 2013). This area is mostly responsive to peripheral representation (>= 30°) and lacks the central part of visual field. Our optic flow stimulus spanned a square of 16° x 16° and it is, therefore, likely that it activated V6 and not V6A. Future studies should however include a wide field retinotopic mapping in their procedures to clearly delineate these two regions. A previous fMRI study in human found that responses in the dorsal parieto-occipital sulcus (dPOS, a region that includes V6) were modulated by the vergence angle (Quinlan and Culham, 2007). Our control experiment (see the *‘Control for vergence’* section) ruled out the possibility that our results are affected by vergence. In human, V6 responds to 3D translational egomotion (Sdoia et al., 2009). Its responses to optic flow are also enhanced when the flow is combined with congruent binocular disparity values (Cardin et al., 2012a). These observations and our results suggest that V6 might have a specific role during locomotion. In macaque, V6 is often described as an area that is principally visual. A tracer study showed that anatomically, it is mostly connected to other visual regions, including areas MST and VIP (Galletti, et al., 2001). Its responses are strongly influenced by optic flow signals but are not modulated by inertial motion (Fan et al., 2015). If areas V6 in human and macaque share similar visual properties, like their retinotopic organization (Pitzalis et al., 2006; 2012) or their selectivity to optic flow (Cardin and Smith, 2010; Fan et al., 2015, but see Cottereau et al., 2017), our results suggest that human V6 has a specific role for processing locomotion consistent vestibular inputs. It is, therefore, possible that the homology between human and macaque V6 is not as pronounced as currently believed (Pitzalis et al., 2013; 2015).

Our results also suggest an implication of area VIP in the processing of vestibular inputs,. Responses to Lat GVS in this area did not differ from those measured during the sham condition. In their study, Smith et al. also reported that Lat GVS did not elicit significant responses in this area (Smith et al., 2012). However, our PPI estimation demonstrated that connections between VIP and areas CSv and PIC were significantly stronger during Lat GVS than during baseline (figure 5-A). The results of this connectivity analysis suggests that area VIP might be implicated in the processing of Lat GVS even though further investigation is needed to better understand its exact role in this condition. VIP responses were significantly stronger during AP GVS (figure 3) and VIP was also more connected to the other ROIs during this condition (see the additional connection to hMT+ in figure 5-B). VIP is, therefore, involved during AP GVS and could be included in a cortical network processing vestibular signals, with a strong preference for the antero-posterior direction. In human, VIP is activated by different depth cues such as egomotion compatible optic flow (Wall and Smith, 2008), and disparity (Yang et al., 2011). This area is the putative homologous of macaque VIP, see e.g. Bremmer et al. (2001), a multisensory area that integrates visual and vestibular inputs (Bremmer et al., 2002; Schlack et al., 2002; Chen et al., 2011). In particular, VIP in macaque strongly responds to optic flow (Cottereau et al., 2017) and is supposed to play an important role for navigation in space (Bremmer, 2005). Altogether, these results are in line with our findings and suggest that area VIP is important for locomotion in both human and macaque.

Significant activations were found in both hMT+ and CSv during AP and Lat GVS conditions. For lateral stimulation, our results are consistent with those of Smith et al., (2012). This study found that CSv had the strongest responses for this condition. This is also the case in our results (see figure 3). Smith et al., also found that MST but not MT was activated during lateral GVS. In our study, we did not perform the localizers that permit to dissociate between MT and MST and we, therefore, only localized the human middle temporal complex (i.e., hMT+) using a functional localizer based on optic flow (see the *‘Materials and methods’* section). The hMT+ complex includes both MT and MST, and might also contain other regions such as the putative homologues of macaque areas FST and V4t (see Kolster, 2010). In our data, we did not find any significant difference between the responses to AP *vs* Lat GVS in both CSv and hMT+. This suggests that the global responses of these areas are equivalent in our two GVS conditions. Note, however, that this does not rule out the possibility that subregions within CSv and/or hMT+ are selective to either one or the other condition. This distinction remains difficult to make at the macroscopic level of fMRI recordings and will need further investigations.

Altogether, the present work supports the idea that, in humans, distinct cortical networks are activated during antero-posterior and lateral GVS.

## Author Contributions

ASC, SC and FAJ contributed to the conception and design of the study and data recording. The experiments and analyses were conceptualized by all the authors. Data analysis was performed by FAJ with substantial input from BRC. All authors were involved in writing the manuscript.

FAJ is now at Institute of Pathophysiology, Johannes Gutenberg University of Mainz.

## Acknowledgments

The authors would like to thank Maxime Rosito for the development of the galvanic stimulation system and its adaptation to the scanner environment. They thank the INSERM U825 MRI technical platform for the MRI acquisitions and Jean-Pierre Jaffrézou for his thorough revision of the English.

